# *De novo* Transcriptome Analyses Provide Insights into Opsin-based Photoreception in the Lantern shark *Etmopterus spinax*

**DOI:** 10.1101/364992

**Authors:** Jérôme Delroisse, Laurent Duchatelet, Patrick Flammang, Jérôme Mallefet

## Abstract

The velvet belly lantern shark (*Etmopterus spinax*) is a small deep-sea shark commonly found in the Easter Atlantic and the Mediterranean Sea. In this study, paired-end illumina HiSeq^TM^ technology has been employed to analyse transcriptome data from eye and ventral skin tissues of the lanternshark species. About 64 and 49 million Illumina reads were generated from skin and eye tissues respectively. The assembly allowed us to predict 119,749 total unigenes including 94,569 for the skin transcriptome and 94,365 for the eye transcriptome while 74,753 were commonly found in both transcriptomes. Among unigenes, 60,322 sequences were annotated using classical public databases. The assembled and annotated transcriptomes provide a valuable resource for further understanding of the shark biology. We identified potential “light-interacting toolkit” genes including multiple genes related to ocular and extraocular light perception processes such as opsins. In particular, a single rhodopsin gene mRNA and its potentially associated peropsin were only detected in the eye transcriptome confirming a monochromatic vision of the lantern-shark. Conversely, an encephalopsin mRNA was mainly detected in the skin transcriptome. The encephalopsin was immunolocalized in various shark tissues confirming its wide expression in the shark skin and pinpointing a possible functional relation with the photophore, *i.e*. epidermal light organs. We hypothesize that extraocular photoreception might be involved in the bioluminescence control possibly acting on the shutter opening and/or the photocyte activity itself.

## Introduction

Unusual and poorly known organisms populate deep parts of oceans. The velvet belly lantern shark *Etmopterus spinax* (Linnaeus, 1758), a common deep-sea shark occurring along the continental shelf of the Eastern Atlantic Ocean and in the Mediterranean Sea [1, 2], is one of these. *E. spinax* has been recently used as a model species for experimental studies on physiological control of its natural luminescence [3-6] however due to practical limitations they have been poorly investigated at a molecular point of view. Practically, molecular data are almost absent from public databases. “*Etmopterus spinax* NCBI research” only permits to find 328 nucleotide sequences and 28 protein sequences (6th July 2018) mainly limited to mitochondrial sequences used for phylogenetic analyses while functional molecular data remains totally missing.

Over the past 450 million years, cartilaginous fish have evolved to fill a large range of predatory niches in marine and freshwater ecosystems [7-9]. The development of a sophisticated battery of sensory systems is considered as an important factor explaining the evolutionary success of the elasmobranchs and their relatives [7]. Sharks have been considered as “swimming noses” because of their high olfactory abilities. Their large telencephalon, *i.e*. the forebrain, is indeed primarily dedicated to olfaction [10, 11]. Other sensory systems - including light perception - received traditionally much less attention [12, 13]. Early studies reported that the retinas of the majority of cartilaginous fishes contained only rod photoreceptors [14, 15] and these organisms were thought to have poor visual acuity with eyes that are specialized for scotopic, *i.e*. dim light, vision with no capacity for photopic, *i.e*. bright light, vision or color discrimination [10]. Rods indeed serve scotopic vision and are highly sensitive, at the expense of visual acuity. Other specialisations include the presence of a tapetum at the rear of the eye for reflecting light back on to the photoreceptors and a high photoreceptor to ganglion cell summation ratio that increases sensitivity at the expense of acuity [14]. More recently, it was demonstrated that the large majority of cartilaginous fishes possess a duplex retina containing both rod and cone photoreceptors [12, 16-20]. Cones are used for photopic and colour vision and are responsible for higher visual acuity. Photoreceptors contain visual pigments, the so-called opsins which are linked to a chromophore prosthetic group related either to vitamin A1 (rhodopsins and chrysopsins) or A2 (porphyropsins), which change its conformation when exposed to light, inducing a cascade that finally transmits the visual information to the brain [12]. According to their environment, different combinations of these pigments are found in sharks: most species, mainly epipelagic species, possess rhodopsins (sensitive to blue green light) while most deep-water sharks have chrysopsins (sensitive to deep blue light), and porphyropsins (sensitive to yellow-red light) are found in some freshwater species [10, 21]. Even if many species are able to function under a range of photopic and scotopic light intensities, some deep-sea shark (*e.g*., *Etmopterus spinax*) and rajids appear to have all-rod retinas [22-25].

Parallel to the visual system, photoreceptor cells can also be involved in non-image-forming light detection. The research on extraocular photoreception was pioneered by Steven and Millott [26-28]. The diffuse photosensitivity over the whole or parts of the animal’s skin was described as the “dermal light sense” and even deeper tissues of the body, such as neural or brain cells, can be photosensitive [26-30]. The photoreceptors present outside the eyes are referred to as extraocular or extraretinal [31, 32]. Like the visual photopigments, non-visual pigments consist of an opsin protein linked to a retinal chromophore. Extraocular photoreception can play an important role in the behavior and the physiology of animals [32]. In sharks, extraocular photoreceptors are commonly known to be associated to the pineal gland [33].

The genome of the Elephant shark, *Callorhinchus milii*, has been sequenced [34, 35] and encodes for four visual opsins: a visual rhodopsin (RHO1) and three color visual opsins (middle wavelength-sensitive, RHO2; long wavelength-sensitive, LWS1 and LWS2) [36]. Unusually, therefore, for a deep-sea fish, the elephant shark possesses cone pigments and the potential for trichromacy. More surprisingly, the genome also encode for 13 non-visual opsins: a pinopsin, a parapinopsin, a RGR-opsin, two TMT-opsins (*i.e., teleost multiple tissue opsin*), a VA-opsin (*i.e., vertebrate-ancient opsin*), an encephalopsin (also designated as panopsin), a peropsin, three neuropsins and two melanopsins (*i.e, non-visual rhabdomeric opsin*) [36].

Here, we report the first transcriptome data of the velvet lantern shark *E. spinax*. De *novo* RNA sequencing was performed on the tapeta-equipped eye containing the all-rod retina [37] and on ventral integument tissues of the shark, *i.e*. main light emitting area of the shark. *E. spinax* is indeed able to emit a blue-green ventral glow (λmax=486nm) thanks to thousands of tiny photophores spread in the ventral epidermis [38, 39]. Photophores are composed of a photogenic cell cluster, *i.e*. the photocytes, enclosed in a pigmented sheath and surmounted by a lens. Some pigmented cells playing an iris-like role are located between the lens and the photocytes [38, 39] (**Figure 1**). The aim of this study was to investigate the opsin-based ocular and extraocular photoreception of the lantern shark *E. spinax*. We highlighted multiple actors of the opsin-based phototransduction in ocular and extraocular tissues as well as other “light-interacting actors” [40]. Our results support the idea that the shark receives and integrates constant light information from the environment but also possibly from their own luminous organs. Light reception at the level of a luminous area could be linked to a specific control of the light emission at the level of the photophore as suggested in various other luminous metazoans (*i.e*., *Echinodermata Amphiura, Ctenophora Mneniopsis, Mollusca Sepiola*) [41-46].

**Figure 1.**
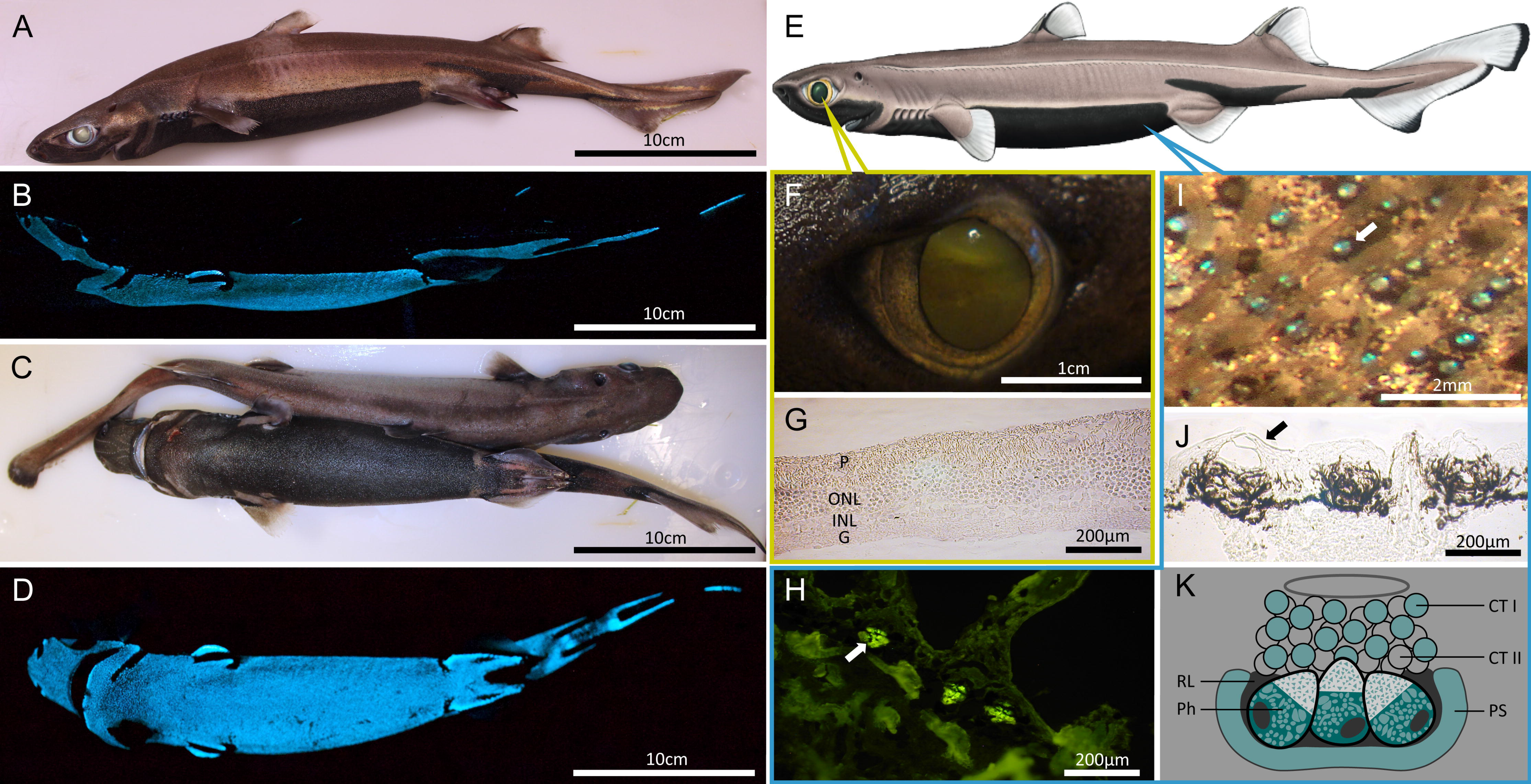
The lanternshark Etmopterus *spinax*. A-B, E: lateral views of the shark (© 2018 Shark Trust, www.sharktrust.org). B: lateral bioluminescence emission pattern. C: oral and aboral view of the shark. C-D: oral views of the shark. F: Eye of the shark. G: histological section of the shark retina. H, J: histological section of the shark skin. K: Schematic reconstruction of a photophore (modified from [38]). I: in vivo observation of ventral skin photophores, J: histological section of the shark retina. Annotations: C: connective tissue, CTI: cellular type I, CTII: cellular type II, D: denticle, L: lens, G: ganglionic layer, E: epidermis, INL: inner nuclear layer, Ir: iris, ONL: outer nuclear layer, P: pigmented layer, Ph: photocyte, PS: pigmented shield, RL: reticulated layer.

## Material and Methods

### Tissue Collection and preparation, Ethics Statement & RNA isolation

Adult velvet belly lantern sharks, *E. spinax* were captured by long-lines lowered at 200m depth in the Raunefjord, Norway (60°169 N; 05°089 E) (see also [38, 39] for more details) during multiple field sessions between August 2014 and January 2016. Living sharks were kept at Bergen University Marine Station (Espegrend, Norway) in a seawater tank (1m^3^) filled with cold (6° C) running seawater pumped from the depths of the adjacent fjord. The tank was placed in a dark room to keep animals under good physiological conditions. Following the local instructions for experimental fish care (Permit 12/14048), captive animals were euthanized by a blow to the head followed by a full incision of the spinal cord at the back of the head. Animals were treated according to the European regulation for handling of animals in research.

The global methodological pipeline of the study is illustrated in the **Figure 1**.Shark tissues from one shark individual were dissected and directly frozen in liquid nitrogen. Pieces of eye and skin tissues were then permeabilized in RNA*later*^TM^-Ice (Life Technologies) during one night at −20°C following the manufacturer’s instructions and then stored at −80°C until RNA extraction or directly processed for RNA extraction. Total RNA was extracted following the Trizol^®^ reagent based method. The quality of the RNA extracts was checked by gel electrophoresis on a 1.2 M TAE agarose gel, and by spectrophotometry using a Nanodrop spectrophotometer (LabTech International). The quality of the RNA was also assessed by size-exclusion chromatography with an Agilent Technologies 2100 Bioanalyzer.

In parallel, patches of ventral and dorsal skin as well as eye of the shark were removed and stored either in 4% paraformaldehyde phosphate buffer saline (PBS) for 12 hours at 4°C. Then, they were kept at 4°C in PBS until use or directly frozen at −80°C without any treatment. Fixed pieces of ventral and dorsal skin (1cm^2^) were used to perform histological and immunohistochemical analyzes while frozen samples were used to perform immunoblots.

### cDNA Library preparation and sequencing

After treatment of the total RNA with DNase I, magnetic beads with Oligo (dT) were used to isolate mRNAs. The purified mRNAs were then mixed with fragmentation buffer and fragmented into small pieces (100-400bp) using divalent cations at 94°C for 5 minutes. Taking these short fragments as templates, random hexamer primers (Illumina Inc., San Diego, CA, USA) were used to synthesize the first-strand cDNA, followed by the synthesis of the second-strand cDNA using RNaseH and DNA polymerase I. End reparation and single nucleotide A (adenine) addition were performed on the synthesized cDNAs. Next, Illumina paired-end adapters were ligated to the ends of these 3’ adenylated short cDNA fragments. To select the proper templates for downstream enrichment, the products of ligation reaction were purified on a 2% agarose gel. The cDNA fragments of about 200bp were excised from the gel. Fifteen rounds of PCR amplification were carried out to enrich the purified cDNA template using PCR primer PE 1.0 and 2.0 (Illumina Inc., San Diego, CA, USA) with phusion DNA polymerase. Finally, the cDNA library was constructed with 200bp insertion fragments. After validation on the Agilent Technologies 2100 Bioanalyzer, the library was sequenced using Illumina HiSeq^TM^ 2000 (Illumina Inc., San Diego, CA, USA), and the workflow was as follows: template hybridization, isothermal amplification, linearization, blocking, sequencing primer hybridization, and first read sequencing. After completion of the first read, the templates are regenerated *in situ* to enable a second read from the opposite end of the fragments. Once the original templates are cleaved and removed, the reverse strands undergo sequencing-by-synthesis. High-throughput sequencing was conducted using the Illumina HiSeq^TM^ 2000 platform to generate 100-bp paired-end reads. cDNA library preparation and sequencing were performed at Beijing Genomics institute (BGI, Hong Kong) according to the manufacturer’s instructions (Illumina, San Diego, CA, USA). After sequencing, raw image data were transformed by base calling into sequence data, which were called raw reads, and stored in the fastq format.

### *De novo* assembly and read mapping

A reference *de novo* transcriptome assembly was performed from *E. spinax* reads derived from eye and skin tissues. Before the transcriptome assembly, the raw sequences were filtered to remove the low-quality reads. The filtration steps were as follows: 1) removal of reads containing only the adaptor sequence; 2) removal of reads containing over 5% of unknown nucleotides ‘‘N’’; and 3) removal of low quality reads (those comprising more than 20% of bases with a quality value lower than 10). The remaining clean reads were used for further analysis. Quality control of reads was accessed by running the FastQC program [47].

Transcriptomed *de novo* assembly was carried out with short paired-end reads using the Trinity software [39] (*version release-20121005; min_contig_length 100, group_pairs_distance 250, path_reinforcement_distance 95, min_kmer_cov 2*). Trinity partitions the sequence data into many individual de Bruijn graphs, each representing the transcriptional complexity at a given gene or locus, and then processes each graph independently to extract full-length splicing isoforms and to tease apart transcripts derived from paralogous genes. After Trinity assembly, the TGI Clustering Tool (TGICL) [48] followed by Phrap assembler (http://www.phrap.org) were used for obtaining distinct sequences. These sequences are defined as unigenes. Unigenes can either form clusters in which the similarity among overlapping sequences is superior to 94%, or singletons that are unique unigenes.

As the length of sequences assembled is a criterion for assembly success, we calculated the size distribution of both contigs and unigenes. To evaluate the depth of coverage, all usable reads were realigned to the unigenes using SOAP aligner with the default settings [49]. Additionally, the completeness of the transcriptomes was evaluated using tBLASTn search against the 248 “Core Eukaryotic Genes” [50].

BLASTx alignments (E-value threshold < 1e^-5^) between unigenes and protein databases like NCBI nr (http://www.ncbi.nlm.nih.gov/), Swiss-Prot (http://www.expasy.ch/sprot/), KEGG (http://www.genome.jp/kegg/) and COG (http://www.ncbi.nlm.nih.gov/COG/) was performed, and the best aligning results were used to identify sequence direction of unigenes. When results from different databases are conflicting, the priority order nr, Swiss-Prot, KEGG, and COG was followed to decide on sequence direction for unigenes. When a unigene was unaligned in all of the above databases, the ESTScan software (v3.0.2) [51] was used to decide on its sequence direction. ESTScan produces a nucleotide sequence (5’–3’) direction and the amino sequence of the predicted coding region.

For both transcriptomes, unigene expression was evaluated using the “Fragments per kilobase of transcript, per million fragments sequenced” (FPKM) method. The FPKM value is calculated following the specific formula *FPKM* = 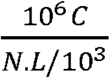where C is the number of fragments showed as uniquely aligned to the concerned unigene, N is the total number of fragments that uniquely align any unigene, and L is the base number in the coding DNA sequence of the concerned unigene. The FPKM method integrates the influence of different gene length and sequencing level on the calculation of gene expression.

The completeness of the transcriptomes was evaluated using tBLASTn search for the 456 human transcripts, from the Core eukaryotic gene dataset, that are highly conserved in a wide range of eukaryotic taxa and has been previously used to assess the quality of genomes and transcriptomes (http://korflab.ucdavis.edu/datasets/cegma/) [50].

### Functional gene annotation of *E. spinax* transcriptome

Following the pipeline described in the **Figure 2**, all unigenes were used for homology searches against various protein databases such as NCBI NR, Swissprot, COG, and KEGG pathway with the BLAST + software (BLASTx, Evalue < 1e^-5^). Best results were selected to annotate the unigenes. When the results from different databases were conflicting, the results from nr database were preferentially selected, followed by Swissprot, KEGG and COG databases. Unigene sequences were also compared to nucleotide databases NT (non-redundant NCBI nucleotide database, E-value < 1e^-05^, BLASTn, http://www.ncbi.nlm.nih.gov).

**Figure 2.**
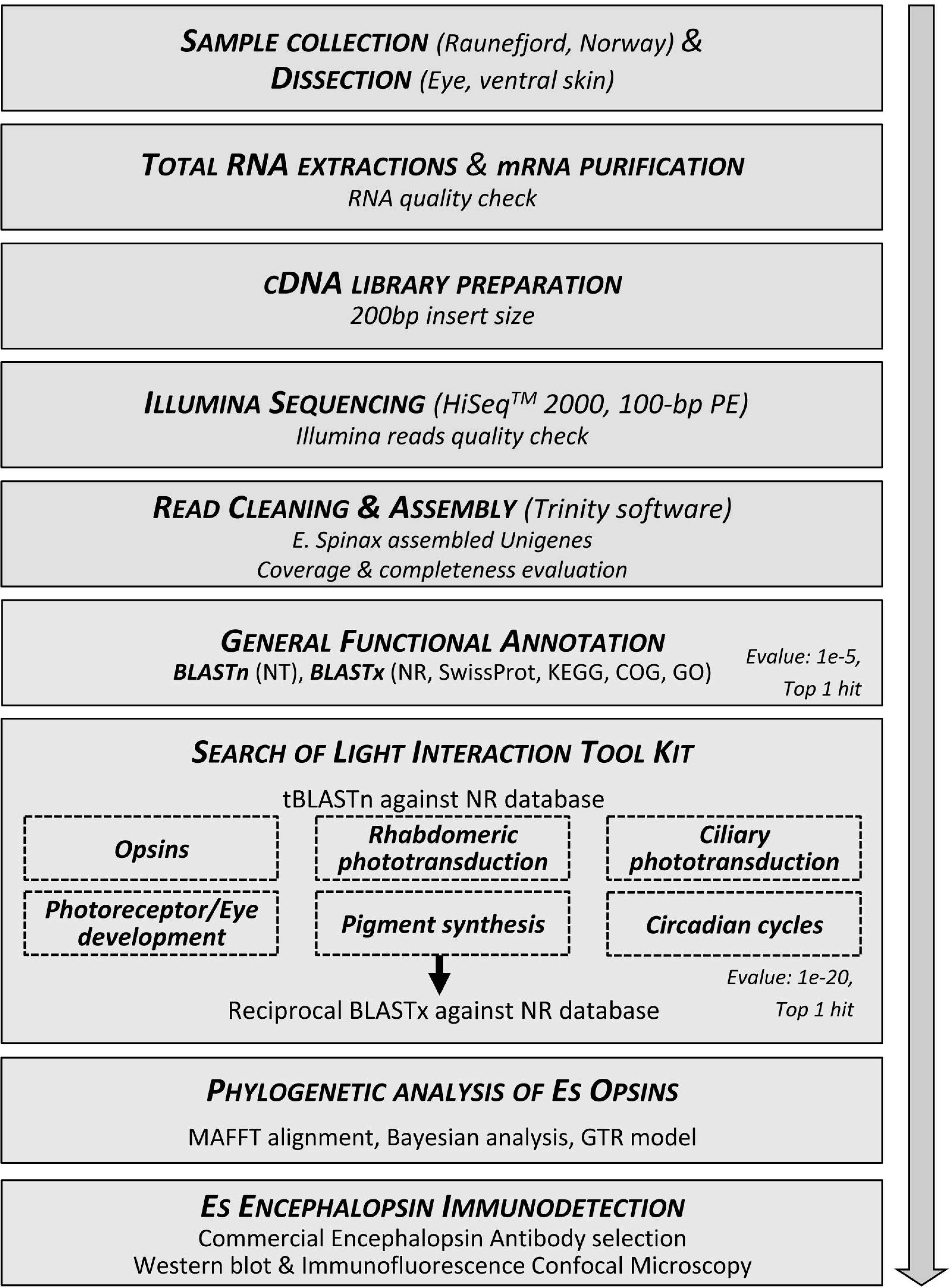
Methodological pipeline of the study performed on the shark *E. spinax*.

To further annotate the unigenes, the Blast2GO program [52] was used with NR annotation to get GO annotation according to molecular function, biological process and cellular component ontologies (http://www.geneontology.org). Gene Ontology (GO) is an international standardized gene functional classification system that offers a dynamically updated and controlled vocabulary and a strictly defined concept to comprehensively describe properties of genes and their products in any organism. After getting GO annotation for every unigenes, the WEGO software [53] was used to establish GO functional classification for all unigenes and to highlight the distribution of gene functions. The unigene sequences were also aligned to the COG database to predict and classify possible functions. COG is a database in which orthologous gene products are classified. Every protein in COG is assumed to have evolved from an ancestor protein, and the whole database is built on coding proteins with complete genome as well as system evolution relationships of bacteria, algae and eukaryotic organisms. Finally pathway assignments were performed according to the KEGG pathway database. The KEGG pathway database records networks of molecular interactions in cells, and variants of them specific to particular organisms.

### Detection of opsins and “light interacting toolkit” genes in *E. spinax*

In order to study genes involved in light-mediated processes such as opsin-based phototransduction (*i.e*., opsins themselves, actors involved in phototransduction associated to rhabdomeric or ciliary opsins), photoreceptor specification, eye development/retinal determination network, retinoid pathway, melanin pigment synthesis, crystallins, diurnal clock and circadian cycles, potential genes of interest were selected based on the phylogenetically-informed annotation (PIA) tool developed to search for light-interacting genes in transcriptomes of non-model organisms [40]. For specific opsin searches, the PIA dataset was implemented with various reference metazoan opsins based on [54] to cover the whole opsin diversity. First, the “Light Interaction Genes” were searched in the transcriptomes of *E. spinax* using *BLAST* analyses (1 hit, Evalue>e-20). All individual unigenes isolated were then reciprocally BLASTed in the NR database (GenBank, RefSeq, EMBL, DDBJ, PDB databases) using Geneious^®^ (v.8.1.9) (tBLASTn with 1 maximum hit) [55]. Reference genes associated with all light-mediated processes are listed in the **Supplementary Table 1**. Blast hits with significant E-values strongly indicate homolog proteins. *E. spinax* homologs should then have BLAST hits to query sequences with significant E-value. In parallel, searches were performed on two chondrichthyan reference genomes: *Rhincodon typus* (22 march 2017; predicted proteins; 27,896 sequences; 13,150,867 total letters) and *Callorhinchus milii* (12 may 2014; predicted proteins; 28,237 sequences; 17,563,624 total letters).

### Opsin characterisations and phylogenetic analyses

For all putative opsin candidates, secondary structure prediction – in particular, of the transmembrane helices – was performed using the MENSAT online tool [56-58]. In *silico* translation (Expasy translate tool, http://expasy.org/tools/dna.html) was performed on the opsin-like sequences retrieved from the *E. spinax* transcriptome. A multiple amino-acid alignment of the putative opsins was performed using Geneious^®^ [55] using MAFFT algorithm [59]. Aligned residues were highlighted by similarity group conservation (defined by the software) and similarity comparisons were calculated in SIAS website platform (http://imed.med.ucm.es/Tools/sias.html). Sequence alignments made it possible to identify opsin characteristic features such as the lysine residue involved in the Schiff base linkage, the counterion, the amino acid triad present in the helix involved in the G protein contact, or putative disulfide bond sites. The predicted molecular weight of the opsins was calculated using the “Compute pI/Mw tool” on the ExPASy Proteomics Server [60].

Phylogenetic analyses of *E. spinax* opsin sequences and reference metazoan opsins, was constructed using trimmed sequence alignment, *i.e*. in particular, the conserved 7TM core of the protein discarding opsin N-terminal and C-terminal sequence extremities to avoid unreliably aligned regions. Automatic alignment trimming was performed using BMGE software (http://mobyle.pasteur.fr/cgi-bin/portal.py) using default parameters. Opsin sequences from other chondrichthyan species, either published or available in online databases, were added in the analysis (**Supplementary table 1**). Sequences of a non-opsin GPCR (*i.e*. melatonin receptor) were chosen as outgroup following previous reference studies [61-63]. Maximum likelihood phylogeny was constructed using PHYML [64, 65]. We performed a Bayesian analysis with MrBayes 3.2 software [66] using the GTR+G model based on recent opsin studies [61-63]. Four independent runs were performed, until a standard deviation value inferior to 0.01 is reached (after 3,500,000 generations).

### Encephalopsin immunodetection on membrane

We used a commercial polyclonal antibody directed against human encephalopsin (anti-H.sapiens encephalopsin Pab, Genetex, GTX 70609, lot number 821400929) to immunolocalised the encephalopsin in *E. spinax. E. spinax* encephalopsin protein sequence, predicted in this study based on RNA-seq data, appears highly similar to other vertebrate orthologous encephalopsins. *E. spinax* predicted encephalopsin protein shares 52% of identity and 61% of similarity with human encephalopsin. To specifically check the antibody specificity, antibodies were tested on membrane after protein extraction, SDS-page and Western blot. Protein extraction of frozen skin samples was performed in two steps: first homogenization with 1000 μl of TEN buffer (10 mM Tris, pH 7,5; 1mM EDTA, pH 8,0; 100mM NaCl) supplemented with protease inhibitors (complete – Mini tablets, Roche). After homogenizing the solution, the sample was sonicated and centrifuged at 800g for 10 min at 4°c. Supernatant was recovered and the remaining solid phase was re-homogenized with 200 μl of TEN buffer containing 10% NP-40 and 0,25% SDS (10mM Tris, pH 7,5; 1mM EDTA, pH 8,0; 100mM NaCl; 0,5 % NP-40; 0,25% SDS; 0,5% Deoxycholate) with protease inhibitors. After further mixing, the sample was sonicated and centrifuged (15 min, 100 000g, 4°C). A second supernatant liquid is collected. Whole process takes place in cold (4°C). To standardise the manipulation and use the same amount of protein in each well of the gels used for Western blotting, a BCA assay was performed using Pierce^TM^ BCA Protein Assay Kit (Thermo Scientific).

Laemmli buffer (Biorad) and β-mercaptoethanol (βMSH, Biorad) were added to each protein extract. The proteins were electrophoretically separated at 200 V for 35 min. The separated proteins were then electroblotted on a nitrocellulose membrane. Membrane was incubated overnight with the primary anti-encephalopsin antibody and with secondary antibody (ECL HRP conjugated anti-rabbit antibody, Life Sciences, NA934VS, lot number 4837492) for 1h. Antibody detection was performed with the reagents of the detection kit (HRP Perkin-Elmer, NEL 104) following the manufacturer instructions. The dilution for the primary antibody was 1:2000. In order to determine the specificity of the observed band, control experiments were included: (*i*) omission of the primary antibody; (*ii*) validation of membrane protein extraction and western blot protocols using an anti-cadherin antibody (Purified Mouse Anti-E-Cadherin (BD Transduction Laboratories, 610181). Expression of encephalopsin protein was also assessed in shark’s retina protein extract and section. Encephalopsin is generally considered as being expressed in the brain [67] but not in the eye [68, 69]. More recently, encephalopsin was shown to be expressed within the retina as well as in a variety of extra-retinal tissues such as brain, testis, liver, lung and skin [70-75].

### Encephalopsin Immunohistofluorescence

Skin patches were bathed in PBS with increasing sucrose concentrations: 10% for 1h, 20% for 2h, and finally 30% sucrose overnight. These tissues were then embedded in O.C.T. compound (Tissue-Tek, The Netherlands) and quickly frozen in isopentane chilled with liquid nitrogen. Thin sections cut with a cryostat microtome (CM3050 S, Leica, Germany) were laid on coated slides (Superfrost, Thermo scientific) and left overnight to dry. Slides were blocked with TTBS (Trizma base (Sigma) 20mM, NaCl 150mM, pH 7,5 + 1% Tween 20 (Sigma)) containing 5% BSA (Amresco). Sections were incubated overnight at 4°C with the commercial encephalopsin antibody used at a dilution of 1:400 in TTBS 5% BSA. Revelation of encephalopsin immune reactivity was done with 1h incubation at RT of fluorescent dye labeled secondary antibody (Goat Anti-Rabbit, Alexa Fluor^®^ 594, Life Technologies Limited) dilution 1:200 in TTBS 5% BSA. In order to label the nucleus of each cell, sections have been subject to a DAPI (DAPI nucleic acid stain, Invitrogen) staining during 15min before being mounted (Mowiol^®^ 4-88, Sigma). Sections were examined using an epifluorescence microscope (Polyvar SC microscope, Leica Reichter Jung) equipped with a Nikon DS-U1 digital camera coupled with NIS-elements FW software. Control sections were incubated in TTBS 5 % BSA with no primary antibody.

## Results and Discussion

### Illumina transcriptome sequencing and *de novo* assembly

In order to characterize the eye and ventral skin transcriptomes of the lantern shark *E. spinax*, mRNA were extracted from frozen tissues. cDNA libraries were generated from isolated mRNA using Illumina HiSeq^TM^ pair-end sequencing technology. 49,178,512 and 64,000,000 raw reads, with the length of 100bp, were generated from a 200bp insert library from eye and ventral skin respectively. Dataset qualities were checked using FastQC software. After low quality reads filtering, the remaining high quality reads (*i.e*., 46,012,442 for retina transcriptome and 51,160,110 for ventral skin transcriptome) were used to assemble the eye and ventral skin transcriptomes with the Trinity software [76]. According to the overlapping information of high-quality reads, contigs were generated. For eye transcriptomic data, the average contig length was 291 pb and the N50 (*i.e*., the median contig size) was of 545 pb. For ventral skin transcriptomic data, the average contig length was 227 pb and the N50 was of 316 pb. Q20 percentages (base quality more than 20) were superior to 95% for both datasets. The GC percentage is around 47% for both transcriptomes. The datasets of raw reads were deposited in NCBI database under Biosample SAMN06293581 accession number.

Using paired-end joining and gap filling, contigs were further assembled into 94,365 unigenes, *i.e*. sequences non redundant unique sequences, for the eye dataset and 93,569 for the ventral skin dataset with a total of 119,749 different unigenes. Eye transcriptome unigenes include 23,183 clusters and 71,182 singletons. Ventral skin transcriptome unigenes contain 14,811 clusters and 78,758 singletons. The size distributions of contigs and unigenes are shown in supplementary material (**Supplementary Figure 1**) and numerical data are summarized in **Tables 1** and **2**.

**Table 1.**
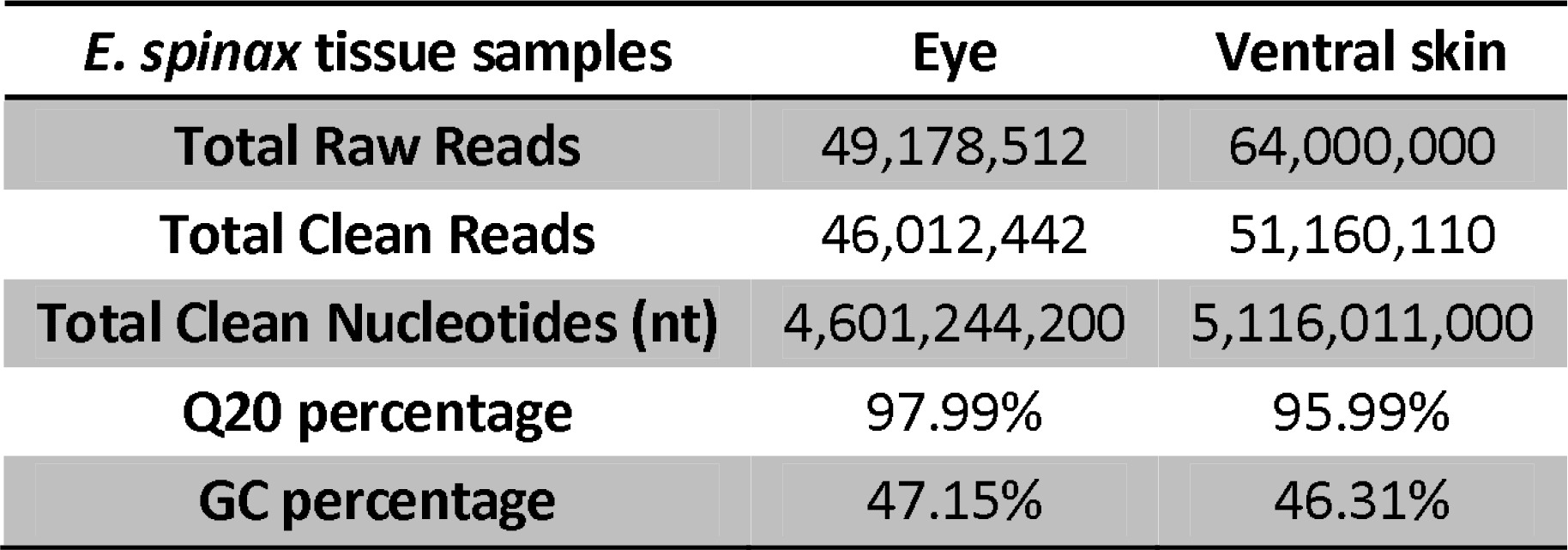
Description of the output sequenced data. Q20 percentage is the proportion of nucleotides with quality value larger than 20 in reads. GC percentage is the proportion of guanidine and cytosine nucleotides among total nucleotides.

| *E. spinax* tissue samples | Eye | Ventral skin |
| --- | --- | --- |
| Total Raw Reads | 49,178,512 | 64,000,000 |
| Total Clean Reads | 46,012,442 | 51,160,110 |
| Total Clean Nucleotides (nt) | 4,601,244,200 | 5,116,011,000 |
| Q20 percentage | 97.99% | 95.99% |
| GC percentage | 47.15% | 46.31% |

**Table 2.**
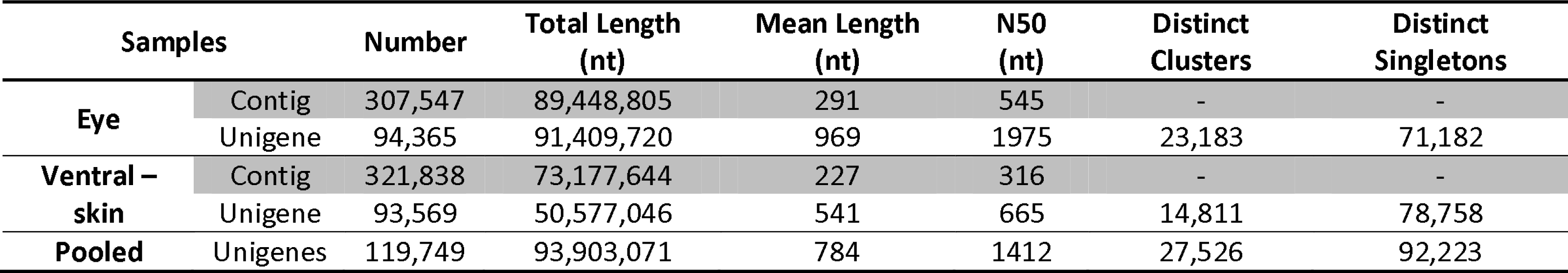
Summary statistics of assemblies for *E. spinax* eye and ventral skin transcriptomes.

The completeness of the transcriptomes was evaluated using tBLASTn search for human transcripts, from the Core eukaryotic gene dataset [50]. A total of 451 (98,9%) of the highly conserved 456 CEGs were detected in the *E. spinax* pooled transcriptome (E-value<1e^-5^). To evaluate the coverage of the two transcriptomes, all the usable sequencing reads were realigned to the all unigenes. More than 78% of eye transcriptome unigenes and more than 76% of ventral skin transcriptome unigenes were realigned by more than 5 reads (**Figure 3**) indicating a good coverage of the whole transcriptome.

**Figure 3.**
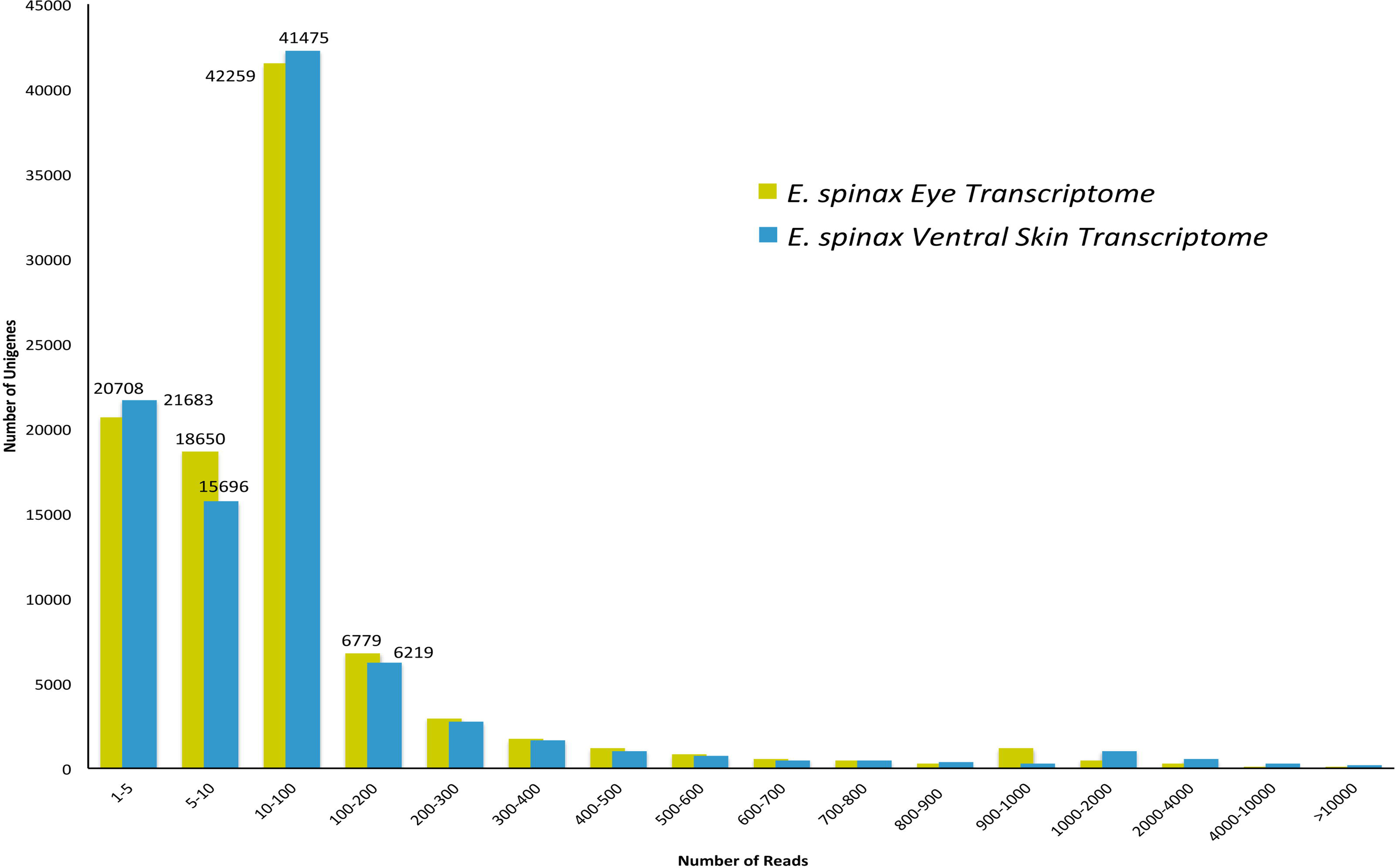
Distribution of the assembled E. spinax unigenes in function of the number of reads to which they can be aligned. The x-axis represents the « number of reads » classes.

Sequence orientations of all unigenes were predicted via ESTScan or BLAST (Basic Local Alignement Search Tool) with an E-value threshold of 10^-5^ in the NCBI database of non-redundant protein (Nr), along with the Swiss-Prot protein database, the Kyoto Encyclopedia of Genes and Genomes (KEGG) and Clusters of Orthologous Groups (COG) database. Finally, total sequence orientation of 56.670 unigenes was predicted for both transcriptomes. On a total of 119,749 predicted unigenes, 20,597 were found in skin transcriptome and 23,077 in eye transcriptome while 73,753 were detected in both transcriptomes. For descriptive purpose, potential DEGs were highlighted, on **Figure 4**, by mapping FPKM values (*i.e*., log_10_(FPKM value ventral skin transcriptome) against log_10_(FPKM value eye transcriptome), calculated for all predicted unigenes. However it has to be clarified that the transcriptome data have been generated in the purpose of new gene discovery, not differential expression analyses, as no biological or technical replication was performed as a part of the study.

**Figure 4.**
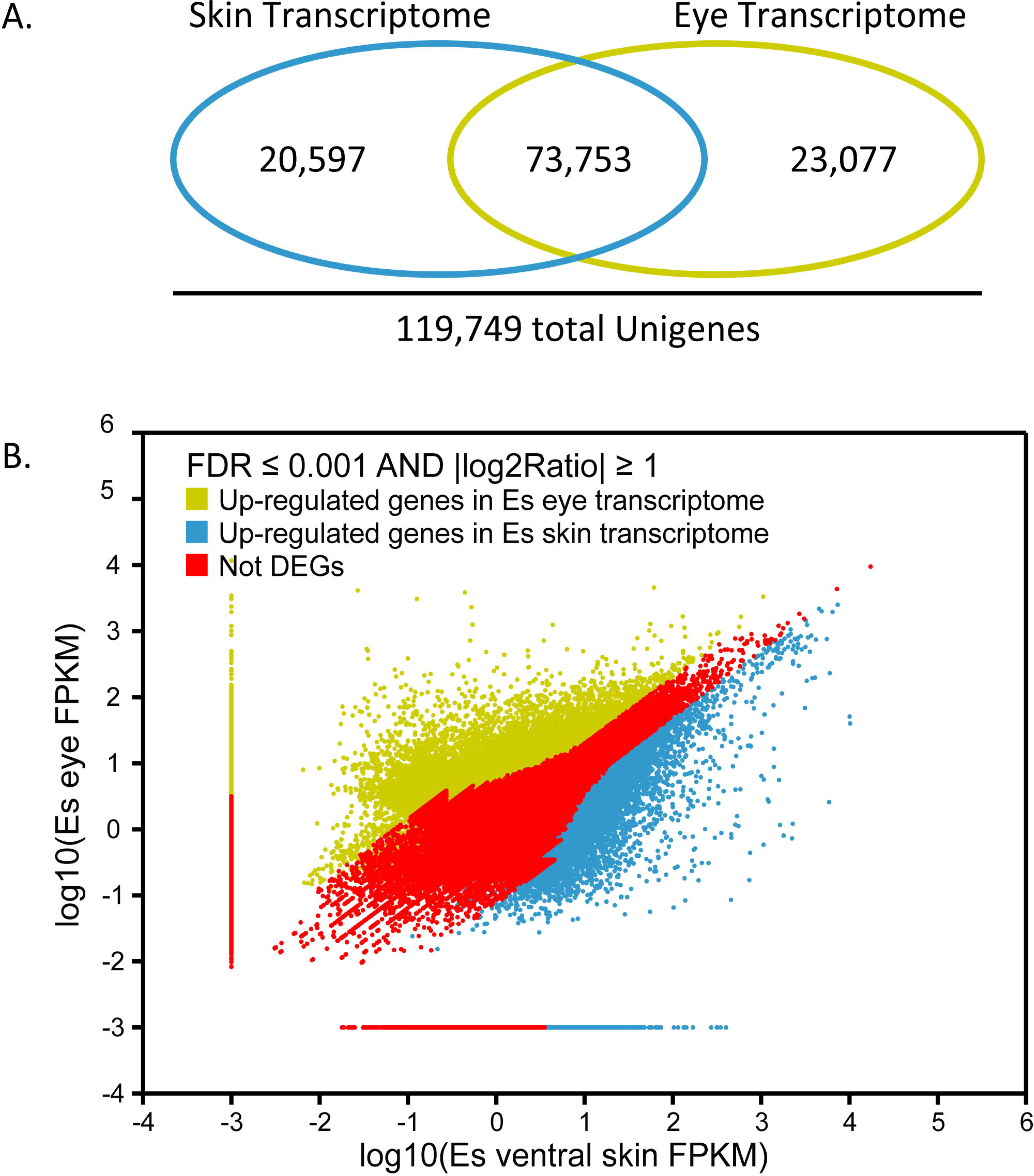
DEGs in *E. spinax* eye and ventral skin tissues.

### Function annotation of *E. spinax* transcriptome

Unigenes sequences were annotated using BLASTx to NCBI protein databases (NR), Swiss-Prot, KEGG and COG with a cut-off E-value of 1e^-5^. Unigenes were also annotated using BLASTn to NCBI nucleotide databases (NT) with a cut-off E-value of 1e^-05^. On all the 119,749 *E. spinax* unigenes, 60,322 show significant matches (50.4%): 46,497 to NR (38.8%), 39,059 to NT (32.6%), 40,078 to Swiss-Prot (33.5%), 34,102 to KEGG (28.5%), 21,768 to COG (18.2%) and 27,753 to GO (23.2%). Because of the lack of genome reference in *E. spinax* and, possibly, the relatively short length of some unigene sequences 49,6% of the assembled sequences could not be matched to any known genes. Annotation results are summarized in the **Figure 5**.

**Figure 5.**
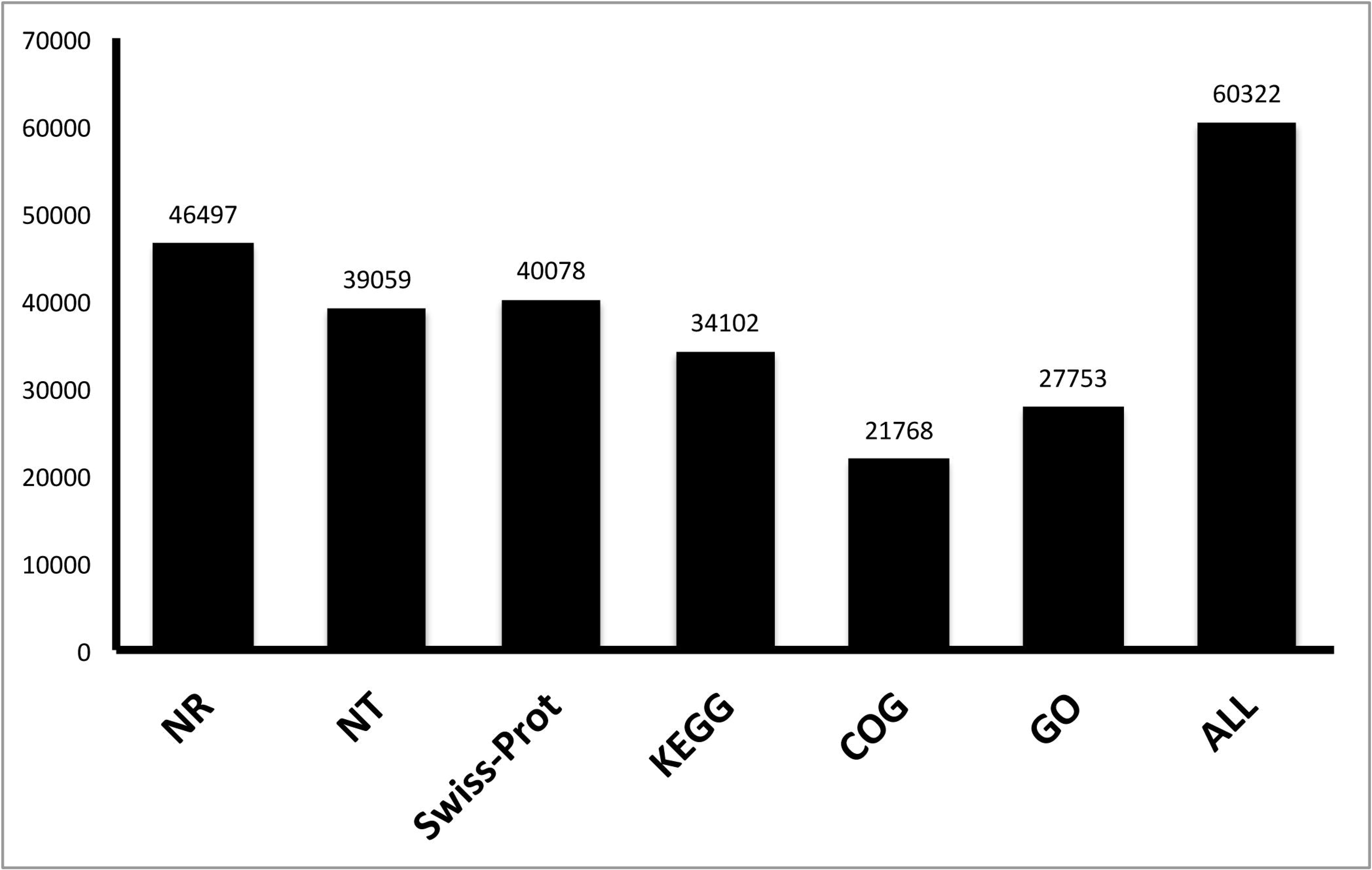
Distribution of annotation results. Unigenes of *E. spinax* were annotated using the nr, nt, Swiss-Prot, KEGG, COG and GO databases (see text for details).

On annotated unigenes from eye and ventral skin transcriptomes, around 8% of sequences were matched to the teleost fish *Silurana tropicalis* (7,8%, **Supplementary Figure 2A**) while 2438 best hits were obtained for the elephant shark *Callorinchus milii*, for which the draft genome is available, 148 for the spiny dogfish *Squalus acanthias* and 122 for the sandy dogfish *Scyliorhinus canicula*. The annotation success was estimated by ranking the annotation E-values and identity results obtained from the NR database comparison. E-value distributions are presented on the **Supplementary Figure 2B**. E-value distribution is highly similar in terms of frequency in both transcriptomes with, for example, around 40% of E-value of « 1e^-15^ to 1e^-5^ » (minimal similarity) in both cases. Similarity distribution is presented on **Supplementary Figure 2C**. For both transcriptomes, around 46% of the mapped sequenced showed significant homology (E value less than 1.0e^-45^) (**Supplementary Figure 2B**) and between 50% of the mapped sequenced have similarity greater than 60% (**Supplementary Figure 2C**).

On the basis of NR annotation, the Blast2go software [52] was used to obtain Gene Ontology annotation of the assembled unigenes, and then the GO functional classifications of the unigenes were performed with WEGO software [53]. For all *E. spinax* unigenes, in total, 27,753 unigenes with BLAST matches to known proteins were assigned to GO classes. Under the category of biological process for example, cellular process (20822; 13% of the biological process total) and single-organism process process (16711; 11% of the biological process total) were prominently represented (**Supplementary Figure 3**). Specific GO categories related to the light perception process, including “Visual perception” (13 hits, GO:0042574), “Phototransduction” (8 hits, GO:0016918), “Retinal binding” (12 hits, GO:0007602) and “Retinal metabolic process” (34 hits, GO:0007601) were targeted in the *E. spinax* pooled transcriptome (data not shown) indicating the expression of phototransduction actors.

FPKM method was used to estimate gene expression in both transcriptome of this study. The 20 most expressed unigenes of eye and ventral skin transcriptomes are shown in the **Supplementary Table 2**. For the eye transcriptome, several actors involved in light perception where highlighted (*e.g*., rhodopsin, Gt protein and crystallins). Within the 20 most expressed unigenes of the ventral skin transcriptome, specific genes include katanin (*i.e*., microtubule-severing protein), keratin and elongation factors. Several common genes, potentially expressed in hematocytes, were highlighted in both transcriptomes (*e.g*., ferritin and hemoglobin). Unsurprisingly, some mitochondrial genes (cytochrome oxidase, NADH dehydrogenase, cytochrome) - linked to eukaryotic energetic metabolism - are highly expressed in both transcriptomes.

### Opsin gene identification, sequence analyses, phylogeny and differential expression

Sequences corresponding to three predicted opsins were found in the *E. spinax* pooled transcriptome. The sequences were translated into protein sequence with Expasy (Expasy, translate tool, Bioinformatics Resource Portal; http://web.expasy.org/translate). Blast analyses revealed they correspond to a rhodopsin, a peropsin and an encephalopsin. These sequences were named accordingly: Es-rhodopsin (complete sequence), Es-peropsin (partial sequence) and Es-encephalopsin (complete sequence). The predicted proteins have molecular weights of 39,654.41 Da, 39,654.41 Da and 46,101.23 Da respectively. Using the MENSAT online tool, characteristic seven transmembrane domains were highlighted in all three sequences. Comparison of the opsin amino acid sequences of *E. spinax* and metazoan opsins demonstrated that the critical residues involved in the maintenance of the tertiary structure of the opsin molecule are present. These key sites include: (*i*) a conserved lysine (K) present in all three Es-opsins and localized at equivalent position 296 of the *H. sapiens* rhodopsin (position 284 for human peropsin, position 299 for human encephalopsin; see **Supplementary Figures 4,5 and 6**) that is covalently linked to the chromophore via a Schiff base [77]; (ii) conserved cysteine (C) residues involved in disulphide bond formation, localized at equivalent positions 110 and 187 for human rhodopsin (98 and 175 for human peropsin, 114 and 188 for human encephalopsin) and present in all Es-opsins [78] which are also conserved throughout the rest of the vertebrate opsin class; (iii) a conserved glutamate (E) at equivalent position 113 of the human rhodopsin that provides the negative counterion to the proton of the Schiff base [79] is also found in Es-rhodopsin; (iv) a conserved glutamate (E) at equivalent position 134 of the human rhodopsin (equivalent position 138 of human encephalopsin) that provides a negative charge to stabilize the inactive opsin molecule [80] is present in Es-rhodopsin and Es-encephalopsin; (vii) the conserved glycosylation sites at equivalent positions 2 and 15 of the human rhodopsin [81] present in Es-rhodopsin (see legends of the **Figure 6 and Supplementary Figures 4, 5, 6** for more details). While there are present in both Rh1 and Rh2 opsins of the elephant shark C. milii, the conserved two cysteine (C) residues at putative palmitoylation equivalent positions 322 and 323 of the human rhodopsin [82] are not conserved in Es-rhodopsin.

**Figure 6.**
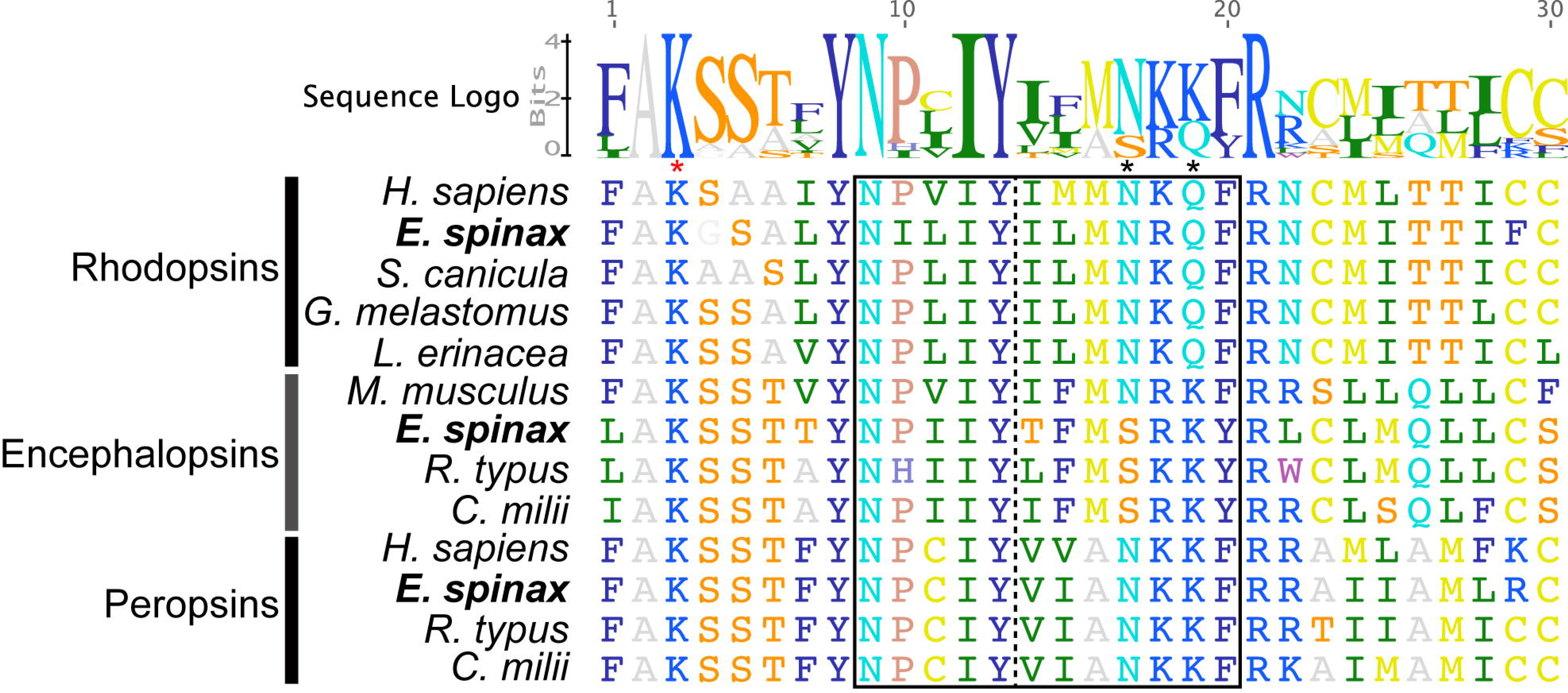
Amino acid alignment of members of three opsin types found in *E. spinax* transcriptomes. *The selected alignment localises to the border (vertical dotted line) between the 7^th^ transmembrane domain and the C-terminal tail. The alignment also includes reference opsins for other metazoans Red asterisk demarcates the position of the Lysine residue critical for Schiff base formation (i.e., K296 of the H. sapiens rhodopsin). The black frame indicated the “NPxxY(x)_6_F” pattern containing the amino acid triad, highlighted with black asterisks (i.e., positions 310-312 in H. sapiens rhodopsin). The “NxQ” motif within the amino acid triad is classically observed in visual c-opsins but is not conserved in encephalopsins. R. typus: Rhincodon typus (encephalopsin: XP_020368171.1, peropsin: XP_020384809), H. sapiens: Homo sapiens (rhodopsin: NP000530.1, peropsin: NP006574.1), L. erinacea: Leucoraja erinacea (rhodopsin: P79863.1), M. musculus: Mus musculus (encephalopsin: AAD32670.1), C. milli: Callorhinchus milli (encephalopsin: XP_007892106.1, peropsin: XP_007895211), G. melastomus: Galeus melastomus (rhodopsin: O93441), S. canicula: Scyliorhinus canicula (rhodopsin: O93459.1), C. conger: Conger conger*.

The sequence of the predicted opsins were then incorporated in a phylogenetic analysis of metazoan opsins. Constructed tree has validated the classification of *E. spinax* predicted opsins into the ciliary opsin group for the Es-rhodopsin (vertebrate visual opsins) and the Es-encephalopsin (vertebrate extraocular opsin, opsin 3 group). Es-Peropsin was also confirmed to belong to peropsin/RGR-opsin group with a clear clustering with vertebrate peropsins. Confidence in this classification is high due to the high posterior probabilities values values (**Figure 7**).

**Figure 7.**
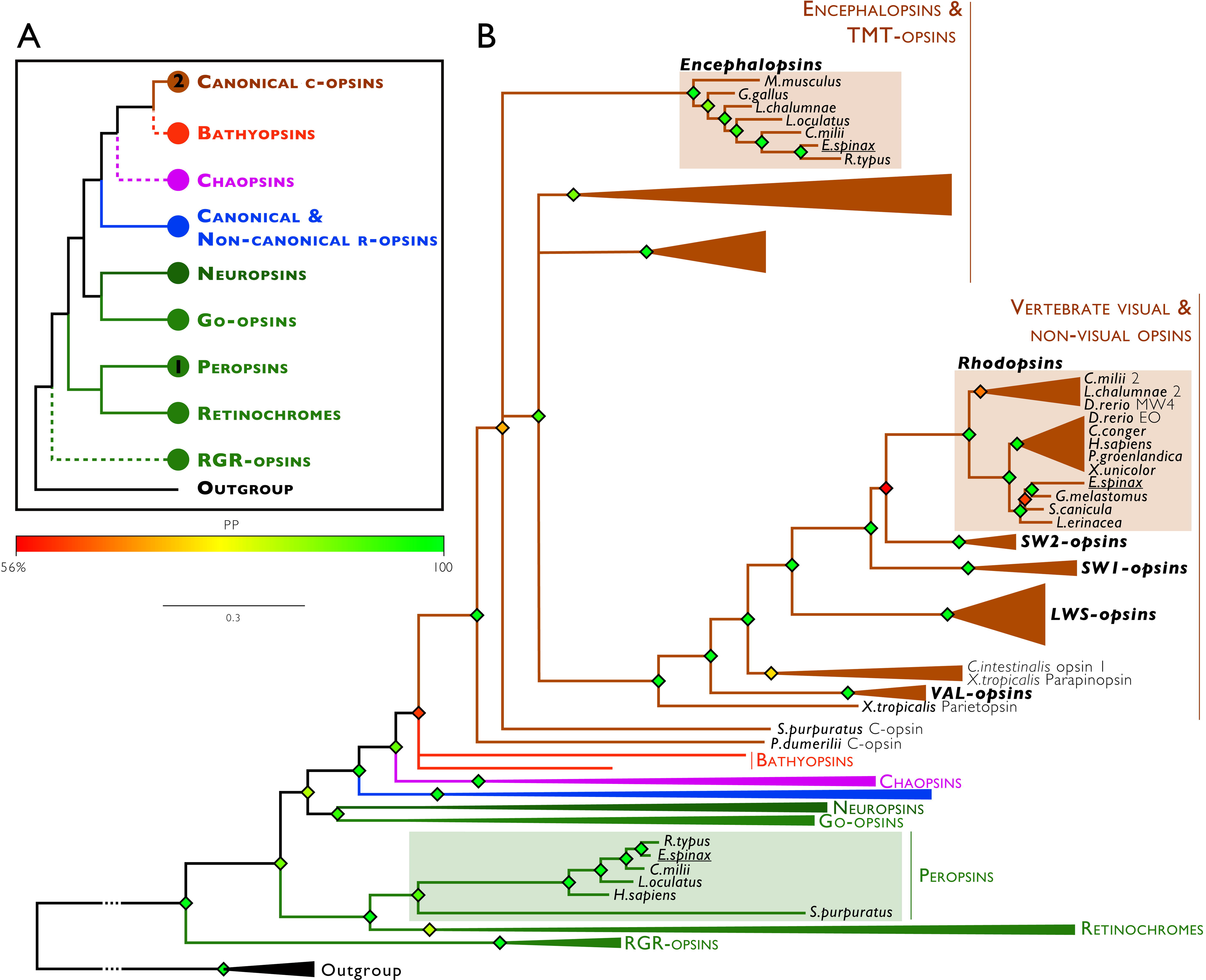
Metazoan opsin phylogenetic tree including the *E. spinax* opsins.

PPredicted E. spinax opsin proteins were included in a large opsin phylogeny (*i*) to ensure their opsin status and (*ii*) define their belonging to known classical opsin groups. Phylogeny was constructed using the Bayesian method (MrBayes software, v.3.2.2). Branch support values are indicated color-codes next to the branching points and correspond to posterior probabilities. Branch length scale bar indicate relative amount of amino acid change. C-opsins: Ciliary opsins, R-opsins: Rhabdomeric opsins, RGR opsin: Retinal G-protein coupled receptors, Outgroup (black): melatonin receptor.

This study does not present a proper differential expression data as no transcriptome replication has been performed. However, differential expression trends can be observed. The Es-rhodopsin and Es-peropsins mRNA were found exclusively in the eye tissue. Based on these observations and on the literature, it seems clear that the rhodopsin and peropsin are functionally coupled as previously described which also confirm the monochromatic vision of the species. Conversely, the Es-encephalopsin was found in both tissues but with a much higher expression in ventral skin (based on FPKM values) (**Figure 8**). Vertebrate encephalopsins belong to the OPN3 group also containing TMT (teleost multiple tissue) opsin in teleosts, pteropsin in insects and c-opsin in annelids [83-85]. OPN3 are non-visual opsins that have been identified in brain of vertebrate and invertebrates [84, 85]. In vertebrate, OPN3 is also expressed in liver, kidney, heart, testes, retina, and epidermis melanocyte and keratinocyte cell types but their function remains unknown [68, 73, 86, 87]. Haltaufderhyde *et al*. [86] suggested that encephalopsin might initiate light–induced signaling pathways contributing to UVR phototransduction in skin. The pufferfish TMT opsin and mosquito OPN3 were shown to have the ability to form a green–light activated photopigment [83]. In Zebrafish TMT-opsin was suggested as a candidate for the photic regulation of peripheral clocks [87]. Concerning all three Es-opsins, the top blast results and the E-value of the hit concerning the reciprocal blast are listed in the **Supplementary Table 1**.

**Figure 8.**
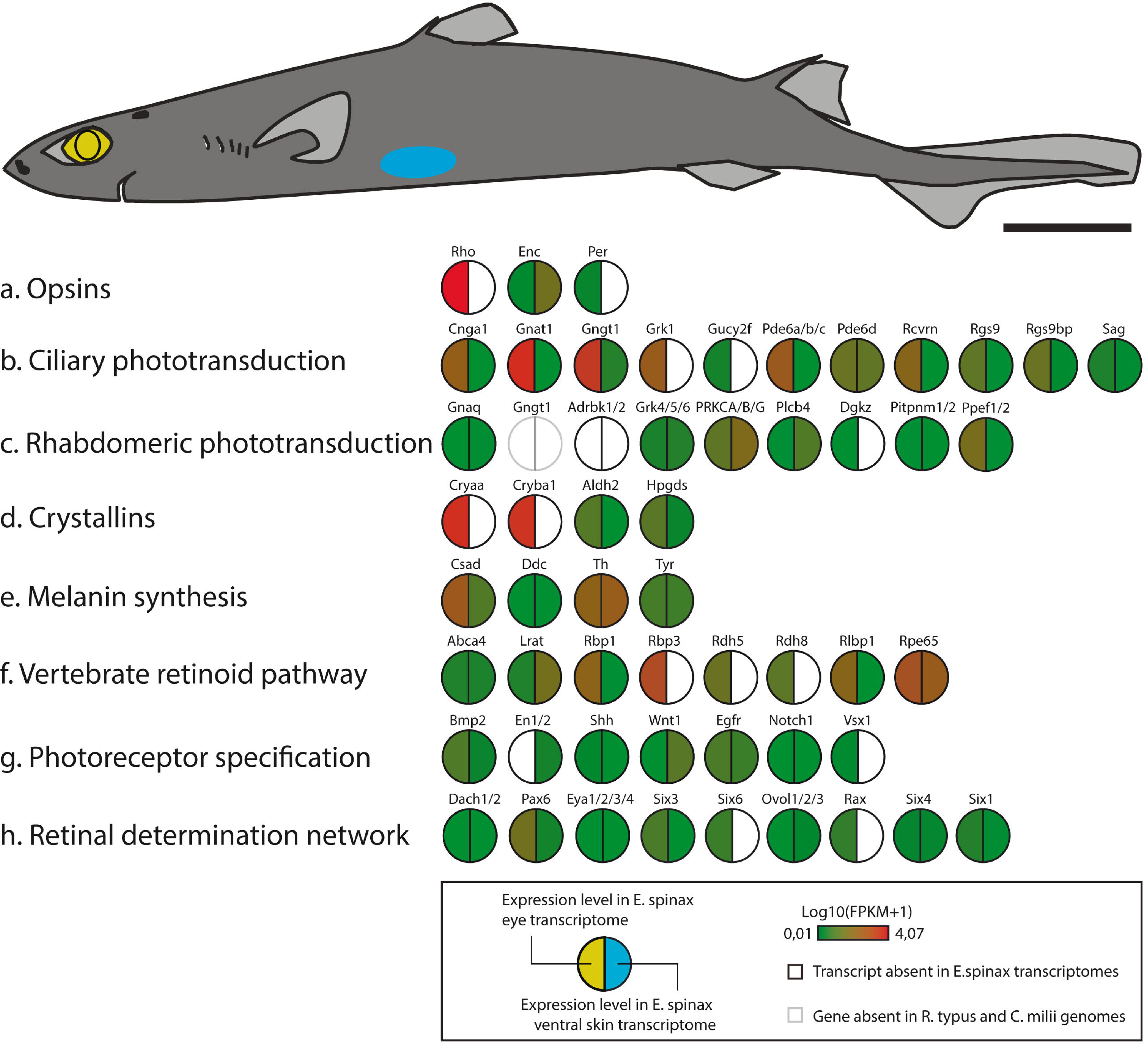
Predicted light-interacting toolkit genes within *E. spinax* eye and ventral skin.

### Phototransduction and “light interacting toolkit” genes identification

Phototransduction is a biochemical process by which the photoreceptor cells generate electrical signals in response to captured photons. Two main phototransduction cascades characterize rhabdomeric and ciliary photoreceptors of metazaons. Ciliary photoreceptors, classically associated with vertebrate eyes, employ a phototransduction cascade that includes ciliary opsins (*e.g*., rho), Gi/Gt (*e.g*., Gnat1) proteins (Go protein-mediated phototransduction cascades were also reported in ciliary visual cells of scallop, amphioxus and lizard parietal eye), phosphodiesterase (*e.g*., Pde6) and cyclic nucleotide gated ion channels (*e.g*., Cnga1). The vertebrate cascade starts with the absorption of photons by the photoreceptive C-opsins. Opsin activation triggers hydrolysis of cGMP by activating a transducing phosphodiesterase 6 cascade, which results in closure of the cGMP-gated cation channels in the plasma membrane and membrane hyperpolarization. The hyperpolarization of the membrane potential of the photoreceptor cell modulates the release of neurotransmitters to downstream cells. Recovery from light involves the deactivation of the light-activated intermediates: photolyzed opsin is phosphorylated by rhodopsin kinase (*e.g*., Grk1) and subsequently capped off by arrestin (*e.g*., Sag); GTP-binding transducin alpha subunit (*e.g*., Gnat1) deactivates through a process that is stimulated by Rgs9. Rhabdomeric photoreceptors, classically associated with invertebrate eyes, employ a cascade involving R-opsins, G protein alpha q (*e.g*., Gnaq), phospholipase C (*e.g*., Plcb4) and transient receptor potential ion channels (TRP, TRPL). Visual signaling is initiated with the activation of R-opsin by light. Upon absorption of a light photon the opsin chromophore is isomerized which induces a structural change that activates the opsin. The photoconversion activates heterotrimeric Gq protein via GTP-GDP exchange, releasing the G alpha q subunit. G alpha q activates the phospholipase C (*e.g*., Plcb4), generating IP3 and DAG from PIP2. DAG may further release polyunsaturated fatty acids (PUFAs) via action of DAG lipase. This reaction leads to the opening of cation-selective channels (*e.g*., TRP) and causes the depolarization of the photoreceptor cells. Ciliary and Rhabdomeric cascades can be deactivated by arrestins and rhodopsin kinases and regenerated by retinal binding proteins [42, 43].

An analysis of the transcriptome database, generated from the eye and ventral skin of *E. spinax*, revealed transcripts encoding proteins with high similarities to the key components of visual transduction cascades. We identified cDNAs including genes encoding for putative visual pigment opsins in both tissues, as well as mRNA coding for protein involved in subsequent activation and deactivation of the cascades. Reciproqual BLASTx analysis demonstrated that some of the predicted “phototransduction actors” are highly similar to the classical phototransduction constituents (**Figure 8**).

Other genes associated to other light related processes, obtained from the Light-Interaction Toolkit (LIT 1.0) [40] were investigated such as crystallins, melanin synthesis actors, vertebrate retinoid pathway actors, photoreceptor specification actors, retinal determination network actors and diurnal clock actors (**Figure 8, Supplementary Table S1**).

### Encephalopsin immunodetection

Based on sequence similarity, a commercial anti-encephalopsin (*H. sapiens*) antibody was selected (Genetex, GTX 70609, lot number 821 400 929). *H. sapiens* encephalopsin have 62% of sequence similarity with the predicted lanternshark encephalopsin sequence (see **Supplementary Figure 6**). Immunoblot analyses allow us to observe a strong immunoreactive band on extract of shark ventral skin tissues using the anti-encephalopsin antibody. The observed labelling corresponds to a protein with a molecular weight of 43kDa matching to the predicted encephalopsin protein (*e.g*., opsins generally have a molecular weight comprised between 39 and 45 kDa [88]. The protein extract from the dorsal skin also shows a similar immunoreactivity (data not shown). Finally, no labelling could be detected in the retina of this shark (data not shown). To confirm that the protocol was efficient for the extraction of transmembrane proteins, controls were also performed with the same extracts of *E. spinax* tissues using an anti-cadherin A (*e.g*., a very abundant protein involved in cell adhesion [89, 90]) antibody (BD Transduction Laboratories, 610181) (data not shown). Negative controls (*i.e*., primary antibody omission) were also performed to confirm the specificity of the secondary antibody (data not shown).

Using immunohistochemistry, a strong anti-encephalopsin immunoreactivity was also observed in the cell membrane of the epidermal cells and the iris-like structure related pigmented cells in *E. spinax* ventral skin sections. Similarly, the cells on the surface of the lens are labelled (**Figure 9B**). *E. spinax* dorsal skin shows a weaker immunoreactivity on cell membranes of the epidermal cells while no staining is observed in the retina (data not shown). Control with omission of the primary antibody did not show any non-specific binding of the secondary antibodies (data not shown).

**Figure 9.**
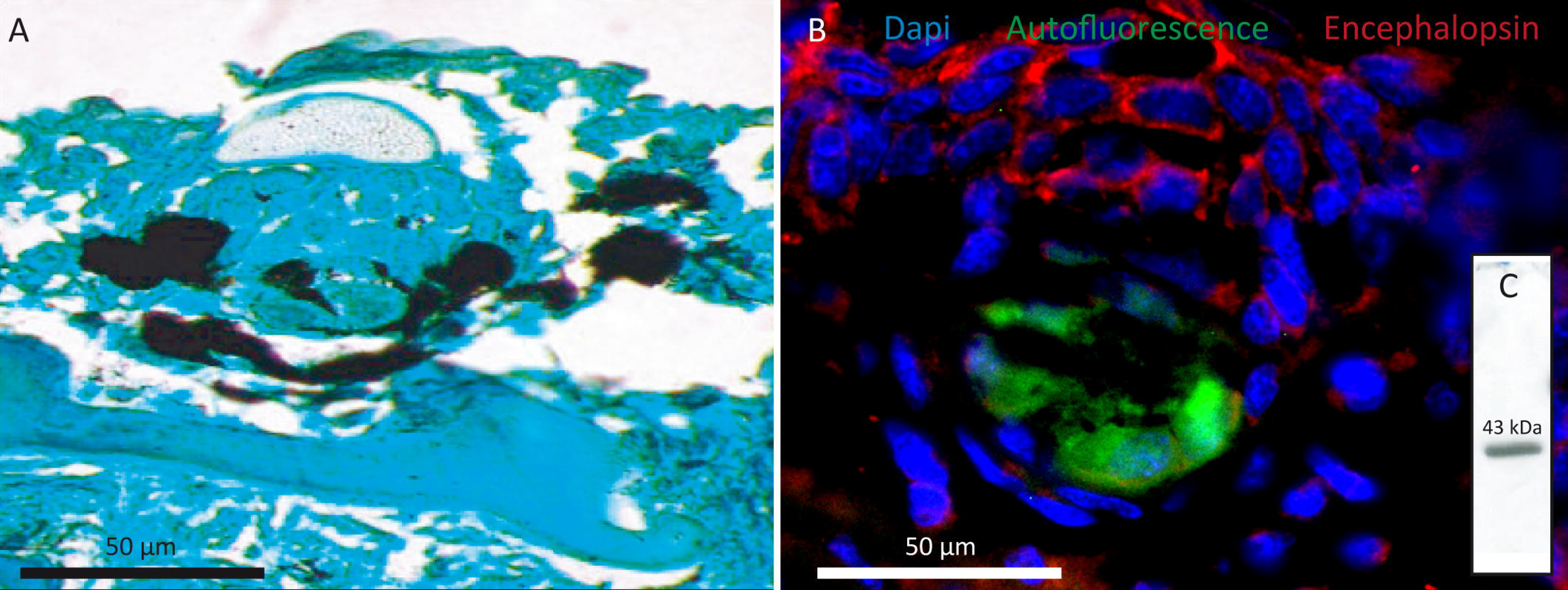
Encephalopsin immunodetection in *E. spinax*. A. photosensitive film of immunoblotting performed on the protein extract of *E. spinax* ventral ad dorsal skin as well as retina with an antibodies directed against extraocular opsin: anti-encephalopsin PAb from Genetex, GTX 70609, lot number 821 400 929, 1/2000. 50ug of total protein were used in each well. B. Cryosection immunofluorescence directed against extraocular opsins in different tissues of the lantern shark, *E. spinax*. Visualization of a photophore paraffin section (A). Visualization of the labelling on cryosections of a ventral skin section with photophores (B), a section of the retina (C). The B and C sections were given the primary antibody GTX (primary antibody: anti-encephalopsin PAb from Genetex, GTX 70609, lot number 821 400 929, 1/50). The secondary antibody was coupled with a red fluorochrome (Alexa Fluor^®^ 594 Goat Anti-Rabbit IgG (H+L) Antibody, highly cross-adsorbed (A-11037), 1/300 from Life Technologies Limited). C, conjunctive tissue; E, epidermis; Ir: iris-like structure related pigmented cell; L, lens cell; Ph: photocyte; Ps: pigmented sheath; D: dermal denticle; R: rod, C: cone layer. Scale bar: 50**μ**m.

## Data availability

Raw reads of eye and ventral skin transcriptomes of *E. spinax*, as well as associated TSA files, will available on NCBI upon publication acceptance (PRJNA369748).

## Conclusion

Next-generation sequencing (NGS) technology allows obtaining a deeper and more complete view of transcriptomes. For non-model or emerging model marine organisms, NGS technologies offer a great opportunity for rapid access to genetic information. The characterization of the *E. spinax* eye transcriptome revealed the presence of the unique visual opsin (Es-rhodopsin) most probably functionally coupled with a peropsin (Es-peropsin). Investigation of ventral skin transcriptome of the lantern shark *E. spinax* revealed the extraocular expression of an encephalopsin, *i.e*. a non-visual ciliary opsin (Es-encephalopsin). Immunodetections of the encephalopsin showed a widespread expression within the cell membrane of the shark epidermis cells surrounding the photophore while no expression is seen in the photocytes themselves. Where darkness is permanent, bioluminescence constitutes the main source of light and these sharks are no exception to the rule. These mid-water cartilaginous fishes indeed emit a ventral light to efficiently mask their silhouette from downwelling ambient light and remain hidden from predator and prey [91]. The encephalopsin expression in the surrounding area of the photophore supports the hypothesis of a potential interaction between light emission and reception. This hypothesis should be confirmed by a deeper characterisation of the *E. spinax* encephalopsin expression and function.

## Acknowledgments

This study was support by a F.N.R.S - F.R.S grant awarded to J.M. and P.F (1887857). Library preparation and sequencing was performed by B.G.I. (Hong Kong). We are grateful to A. Aadnesen, manager of the Espegrend Marine Biological Station (University of Bergen, Norway) where animals were kept and T. Sorlie for his help during field collections.

## Author Contributions

JM and LD collected and prepared the samples during field missions. JD and LD performed the experiments. JD wrote the manuscript. All authors revised the manuscript.

---

**Supplementary Table S1. Search for opsins and “light interacting genes” in the E. spinax eye and ventral skin transcriptomes based on reference sequences.** Homologues to ciliary and rhabdomeric phototransduction components, crystallins, melanin synthesis components, vertebrate retinoid pathway components, photoreceptor specification actors, retinal determination network actors, invertebrate retinoid pathway and diurnal clock components and their reciprocal best BLAST hit in E. spinax transcriptomes. BLAST analyses were also performed on *Rhyncodon typus* [92] and *Callorhinchus milii* [35] genomes. FPKM values and fold change (log10) used for the Figure 8 are shown.

**Supplementary Table S2. Blastn/x of 10 most expressed Unigenes in ventral skin and eye transcriptomes of E. spinax**. Unigene commonly found in both transcriptome hits are in bold.

**Supplementary Figure S1. Distributions of contigs and unigenes sizes in E. spinax retina and ventral skin transcriptomes**. The length of contigs and unigenes ranged from 200 bp to more than 3,000 bp. Each range is defined as follows: sequences within the range of X are longer than X bp but shorter than Y bp.

**Supplementary Figure S2. (A) E-value distributions, (B) similarity distributions and (C) species distributions of the top BLAST hits for all unigenes from *E. spinax* transcriptomes in the nr database.**

**Supplementary Figure S3. Gene ontology classifications of assembled unigenes *E. spinax*.** The results are summarized in three main categories: Biological process, cellular component and molecular function.

**Supplementary Figure S4. *Amino acid alignment of E. spinax rhodopsin with reference metazoan rhodopsins*.**

**Supplementary Figure S5. *Amino acid alignment of E. spinax peropsin with reference metazoan peropsins*.**

**Supplementary Figure S6. *Amino acid alignment of E. spinax encephalopsin with reference metazoan encephalopsins*.**

